# Astrocytic Regulation of aberrant perineuronal net formation in *Mecp2*-null Neocortex

**DOI:** 10.1101/2025.08.13.670146

**Authors:** Ashis Sinha, Angelica M. Kowalchuk, Nasim Khatibi, Russell T. Matthews, Jessica L. MacDonald

## Abstract

Rett syndrome (RTT), caused by mutations in *MECP2*, is a complex neurological disorder characterized by myriad physiological disruptions, including early closure of the critical period of developmental plasticity and precocious formation of perineuronal nets (PNNs). PNNs are lattice-like substructures of extracellular matrix (ECM) that enwrap specific subpopulations of neurons. PNNs are essential in the modulation of neuronal plasticity and brain maturation, and their enzymatic disruption can partially restore plasticity in adults and improve memory. Although precocious PNN formation is well-established in RTT, little is known of the cellular, molecular, or biochemical underpinnings of their precocious formation, or whether precocious PNN formation is due to cell-autonomous or non-cell-autonomous mechanisms. While PNNs form on subsets of neurons throughout the brain, astrocytes secrete many ECM components that form PNNs, and they play a central role in controlling closure of the critical period. We find that *Mecp2*-null astrocyte conditioned media induces the expression of the key PNN component *Hapln1* and causes enhanced PNN formation on wildtype neurons, suggesting that *Mecp2*-null astrocytes play a key role in the precocious formation of PNNs in RTT. Further, we identify increased expression of HAPLN1 and other PNN / ECM components in the developing *Mecp2*-null cortex, and demonstrate that PNNs are structurally and biochemically mature at an earlier developmental stage. These results provide essential insight into the mechanisms and structure of aberrant PNNs in *Mecp2*-null cortex and identify potential new avenues for targeted rescue or reversal of the precocious closing of the critical period in RTT.

## Introduction

Rett syndrome (RTT), an X-linked progressive neurodevelopmental disorder affecting ∼1 in 10,000 live female births (Chahrour and Zoghbi, 2007), is caused by mutations in the transcriptional regulator *MECP2* (Amir et al., 1999; Chahrour and Zoghbi, 2007). Girls with RTT develop relatively normally for 6-18 months, after which they undergo a period of rapid regression, with loss of motor skills, including purposeful hand movement, deceleration of head growth, and onset of repetitive, autistic behaviors (Rett, 1966; Hagberg et al., 1983). *Mecp2* mutant mouse models recapitulate many aspects of human RTT, with one notable difference being that male *Mecp2* hemizygous null (*Mecp2*-/y; KO) mice survive into early adulthood (Guy et al., 2001) and they have been a commonly studied model (Ribeiro and MacDonald, 2020). Studies from both mouse and human have found that MeCP2 regulates numerous genes and cellular pathways in a tissue- and cell-type specific manner (Samaco et al., 2009; Chao et al., 2010; Lioy et al., 2011; Derecki et al., 2012; Sugino et al., 2014), and loss of MeCP2 function in defined CNS circuits results in distinct RTT phenotypes (Alvarez-Saavedra et al., 2007; Fyffe et al., 2008; Adachi et al., 2009; Ward et al.; Nguyen et al., 2013; Wither et al.; He et al., 2014). This proves a significant challenge to the development of therapeutic treatments, and there is currently no effective treatment or cure for this devasting disorder.

While much research has focused on the role of MeCP2 in distinct neuronal subtypes, disruption of *Mecp2* in astrocytes is sufficient to drive numerous RTT phenotypes, and *Mecp2*-null astrocytes can alter the development and function of wildtype neurons in a cell non-autonomous manner (Ballas et al., 2009; Maezawa et al., 2009; Williams et al., 2014; Rakela et al., 2018; Dong et al., 2020; Caldwell et al., 2022). Astrocyte conditioned media (ACM) from *Mecp2*-null astrocytes leads to reduced neuronal size and complexity of both mouse and human-derived neurons and alters synaptic development (Ballas et al., 2009; Williams et al., 2014; Caldwell et al., 2022), and *Mecp2*-null astrocytes lead to distinct transcriptional changes in wildtype neurons (Albizzati et al., 2024). While some differences in the factors secreted by *Mecp2*-null astrocytes have been identified (Ehinger et al., 2021; Caldwell et al., 2022), how *Mecp2*-null astrocytes alter neuronal development and function and contribute to RTT pathology is incompletely understood.

Astrocytes secrete many extracellular matrix (ECM) components that form perineuronal nets (PNNs) (Carulli et al., 2006; Giamanco and Matthews, 2012), and they play a central role in controlling the closure of the critical period (Ribot et al., 2021). PNNs, unique honeycomb- or cobblestone-like structures on the surface of specific subclasses of neurons surrounding synapses on the cell body, predominantly parvalbumin-positive interneurons in the cortex (Hockfield and McKay, 1983; Celio et al., 1998), form coincidently with closure of the critical period for developmental plasticity in the cortex and they are essential in the modulation of neuronal plasticity and brain maturation (Wang and Fawcett, 2012). Early critical period closure and precocious formation of PNNs have been identified in multiple brain regions in *Mecp2*-mutant mice (Krishnan et al., 2015; Krishnan et al., 2017; Sigal et al., 2019; Lau et al., 2020; Patrizi et al., 2020; Carstens et al., 2021) and postmortem human RTT brains (Belichenko et al., 1997; Carstens et al., 2021), supporting the clinical relevance of the altered PNN formation in *Mecp2*-mutant mice. RTT has, in fact, been posited to be a disorder of early critical period plasticity (Picard and Fagiolini, 2019).

Although precocious PNN formation and altered cortical plasticity are well-established in *Mecp2*-mutant mice, little is known currently about the molecular composition and structure of the aberrant PNNs formed in RTT, which is essential to develop strategies to specifically rescue their formation and elucidate mechanisms by which they disrupt plasticity. Further, whether the precocious PNN formation on neurons is due to cell-autonomous or non-cell-autonomous mechanisms is not known. We tested the hypothesis that *Mecp2*-null astrocytes are deficient in the ability to regulate the timing and structure of PNN development, leading to their precocious and altered formation in RTT. Determining how *Mecp2*-null astrocytes alter PNN development in a cell non-autonomous manner will have broad implications for mechanisms that regulate critical period plasticity, both in wildtype and the *Mecp2* mutant contexts.

## Results

### Precocious formation of PNNs in developing Mecp2 KO neocortex

We first confirmed previous studies demonstrating precocious and enhanced staining of PNNs with Wisteria floribunda agglutinin (WFA) on neurons within the male hemizygous null (*Mecp2-/y;* KO) cortex (Krishnan et al., 2015; Sigal et al., 2019; Patrizi et al., 2020). At postnatal day (P)16, WFA labeling is apparent throughout the *Mecp2*+/y wildtype (WT) somatosensory cortex, with abundant labeling within the barrel cortex (Fig. 1A,C). WFA labeling in the *Mecp2* KO is more extensive throughout the somatosensory cortex with a higher overall intensity, particularly in layer VI of the barrel cortex (Fig. 1B,D). Higher magnification images demonstrate similar overall structure of the PNNs, however (Fig. 1E,F), suggesting that the number of PNNs largely accounts for the increased WFA labeling.

**Figure 1.**
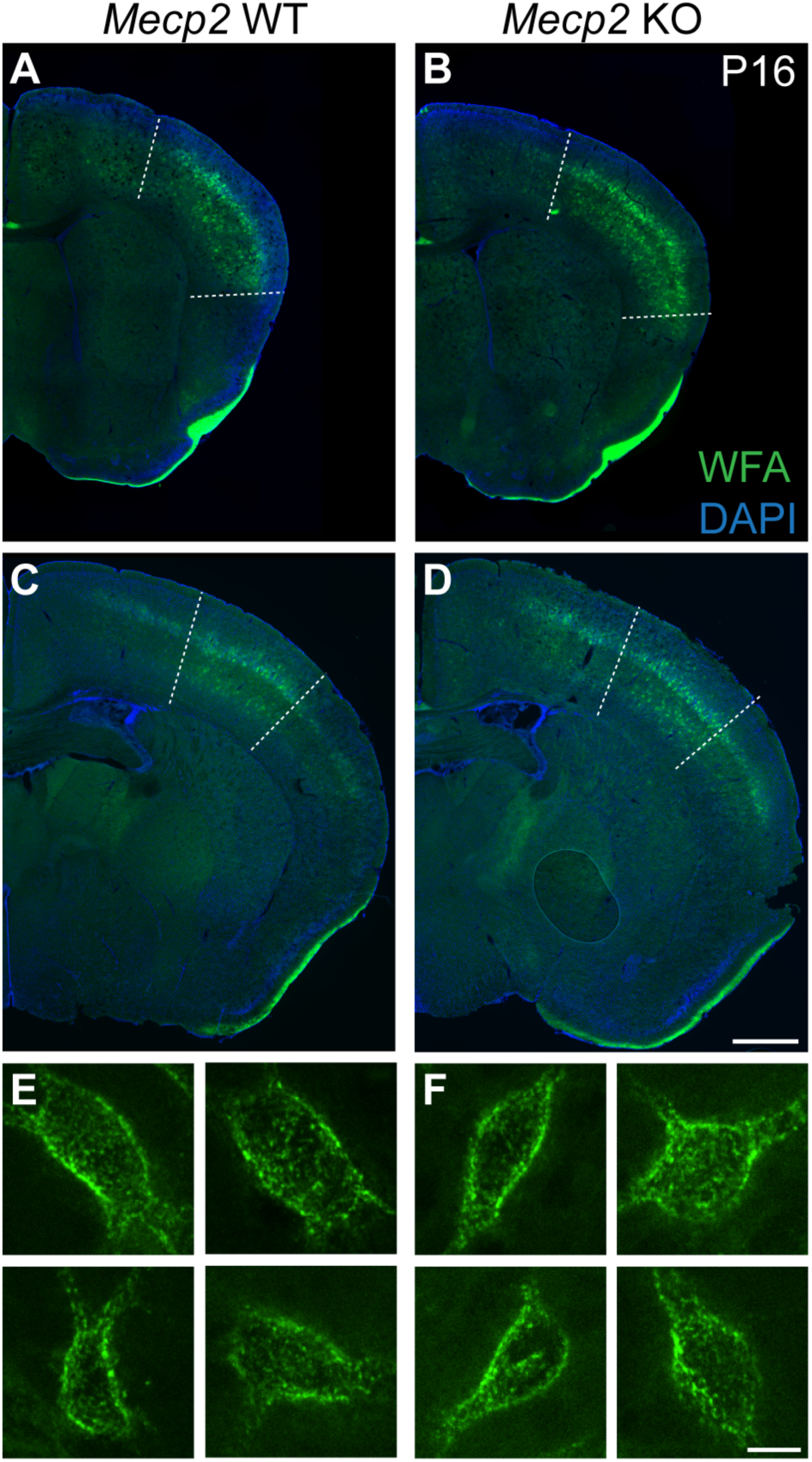
Increased PNN Formation in *Mecp2* KO neocortex compared to *Mecp2* WT at P16. (A-F) WFA labels developing PNNs, particularly evident in the layers IV-IV of the neocortex. More robust and expansive labeling is apparent in the rostral somatosensory region of the KO cortex (B) relative to WT (A) (boundaries indicated by dotted lines). (C,D) This expansion is also seen more caudally down the rostral-caudal axis (barrel cortex boundaries indicated by dotted lines). Scale bar = 500 μm. (A’-B’) High magnification images of WFA+ PNNs do not identify gross morphological changes in PNN structure in the KO (F) compared to the WT (E). High magnification WFA+ PNN images taken from somatosensory cortex in images C and D. Scale bar = 10 μm.

### Astrocytes regulate the timing PNN formation in vitro

The precocious PNN formation in the *Mecp2* KO neocortex could be due to altered developmental trajectory of the PV interneurons on which the PNNs form (Patrizi et al., 2020; Rupert et al., 2023) or altered neuronal activity and interaction between the interneurons and excitatory neurons as PNN formation is activity-driven (Moissidis et al., 2025). In addition to neurons, astrocytes play a role in controlling the closure of the critical period of plasticity (Ribot et al., 2021). *Mecp2* KO astrocytes disrupt neuronal development (Ballas et al., 2009; Maezawa et al., 2009; Williams et al., 2014; Rakela et al., 2018; Dong et al., 2020; Caldwell et al., 2022), raising the question of whether altered PNN development in RTT is due to aberrant astrocytic regulation. To investigate the complex interplay between astrocytes and neurons controlling the development of PNNs, we first examined the formation of PNNs on wildtype primary cortical neurons *in vitro,* in the presence of astrocytes in comparison to when astrocytes are inhibited (Fig. 2). Our previous work indicated that the presence of astrocytes in neuronal cultures delays the formation of aggrecan-expressing ECM accumulation on neurons (Giamanco and Matthews, 2012). Aggrecan is made primarily by neurons and is a key activity dependent component of PNNs (Matthews et al., 2002; Giamanco and Matthews, 2012; Rowlands et al., 2018), suggesting that astrocytes regulate the precise timing of PNN formation.

**Figure 2.**
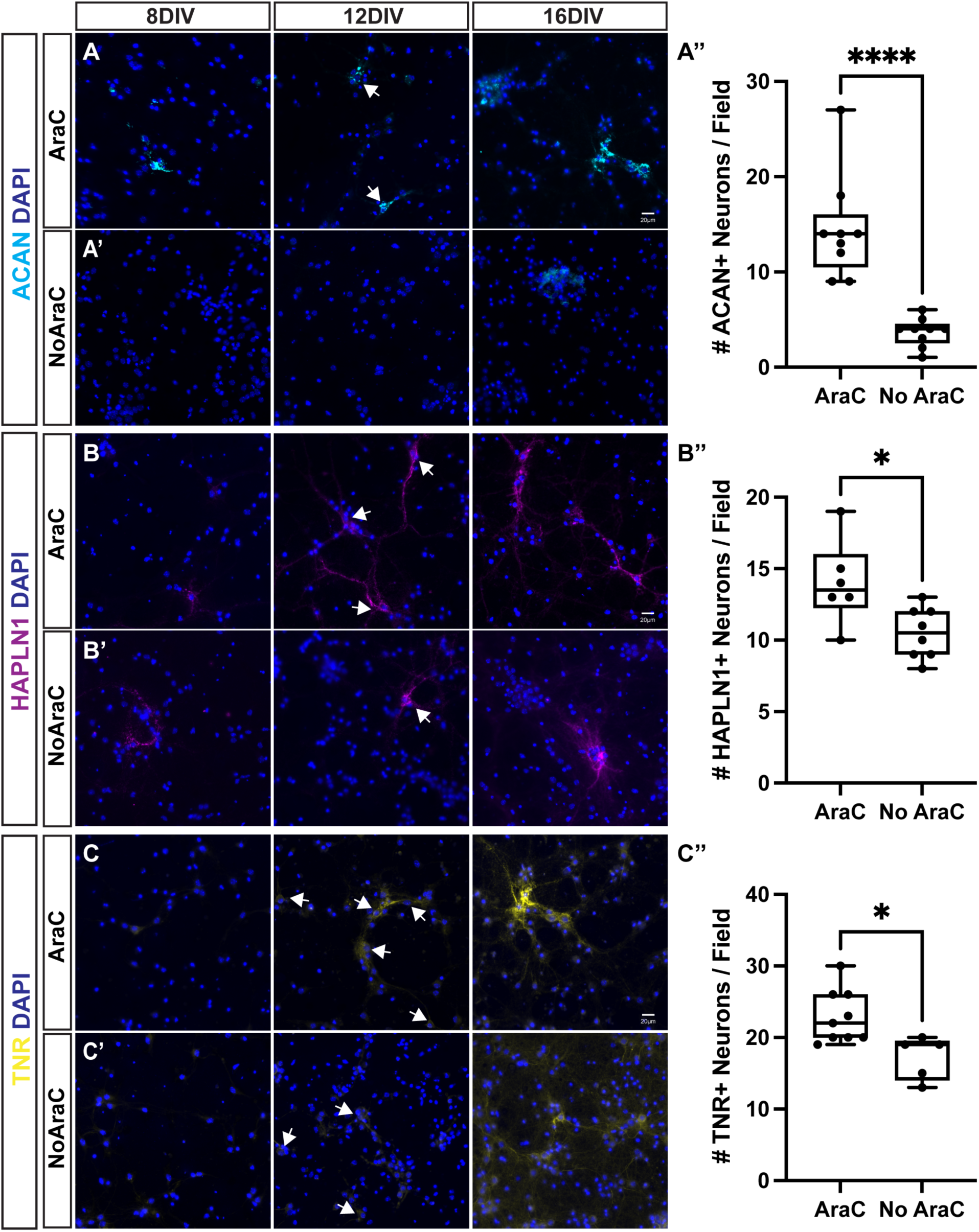
Astrocytes regulate the timing of PNN formation on wildtype neurons *in vitro*. Primary cortical cultures were derived from wildtype CD-1 E15.5 embryos. Cells were either treated with AraC (A,B,C) to deplete glia or left untreated to allow glia to proliferate (NoAraC; A’,B’,C’). Immunohistochemistry was performed for ECM / PNN markers ACAN (A), HAPLN1 (B) and TNR (C) at DIV8, 12 and 16. (A, A’) ACAN+ PNNs first appear around DIV 8 in AraC treated cultures (A) and increase in number and intensity up to DIV 16. In contrast, control NoAraC (A’) cultures had very few ACAN + PNNs even at DIV16. (A’’) The number of ACAN + PNNs (indicated with arrows) was quantified at DIV12, identifying a significant increase in ACAN+ PNNs in AraC treated cells (****, p < 0.0001). (B,B’) As with ACAN, HAPLN1+ PNNs could be observed in the AraC treated cells at DIV8. Compared to this group, HAPLN1 + PNNs were barely observable in control NoAraC cells. (B’’) There was a significant increase in HAPLN1+ PNNs in AraC treated cultures at 12 DIV, as observed with ACAN (*, p = 0.017). (C,C’) Overall TNR detection is greater in control NoAraC cells compared to the AraC group at all time points observed. (C-C’’) However, incorporation of TNR into PNNs is significantly greater in AraC treated cultures at DIV12 (*, p = 0.013). N = 3 - 4 large scans per culture per condition, 3 independent cultures. Bars represent Mean ± S.D. Scale bar 200μm.

To test the developmental timing of PNN formation, astrocytes in E15.5 wildtype cortical cultures were either allowed to proliferate, or they were inhibited by the mitotic inhibitor AraC from 1-3 days *in vitro* (DIV) in parallel experiments. We confirmed by immunohistochemistry that AraC treatment led to a significant reduction in the number of GFAP-positive astrocytes (data not shown). We examined the time-course of expression of key PNN components, as well as their aggregation on the surface of subsets of neuronal cells (Fig. 2). At 6 DIV, PNN / ECM components were not readily detected in either condition (data not shown). Starting by 8 DIV, PNN / ECM components were detectable in AraC treated cultures (Fig. 2A-C) but not non-AraC cultures (Fig. 2A’-C’). By 12 DIV, there were significantly more WT neurons expressing the PNN / ECM components ACAN (Fig. 2A’’), HAPLN1 (Fig. 2B’’), and TNR (Fig. 2C’’) in cultures in which glial cells were inhibited, in comparison to those not treated with AraC. These data suggest that astrocytes play a key role in the developmental timing of PNN formation on neurons, likely preventing precocious formation of the PNNs.

### Mecp2 KO ACM induces unique transcriptional changes in wildtype neurons

How astrocytes regulate the developmental timing of PNN formation on neurons is not known. One possibility is that this phenotype is not the direct result of availability of PNN components from the astrocytes but involves differential activation of signaling cascades in neurons by astrocytes. Unfortunately, the signaling pathways that are key in regulating PNN formation are poorly understood; we thus performed RNA-seq on wildtype neurons treated with astrocyte conditioned media (ACM) in comparison to control non-conditioned media (NCM). Based on the timing of PNN formation observed in these cultures, with differences noted with the presence of astrocytes between 8-12 DIV (Fig. 2), we treated the neurons with ACM starting at 6 DIV and performed bulk RNA-seq on the wildtype cortical neurons at 12 DIV (Fig. 3A). Prior studies have demonstrated that conditioned media from *Mecp2* KO astrocytes is sufficient to significantly disrupt the development of wildtype neurons (Ballas et al., 2009; Williams et al., 2014; Caldwell et al., 2022) thus we investigated whether *Mecp2* KO ACM induces altered transcriptional responses in neurons that could contribute to the disrupted developmental timing of PNN formation.

**Figure 3.**
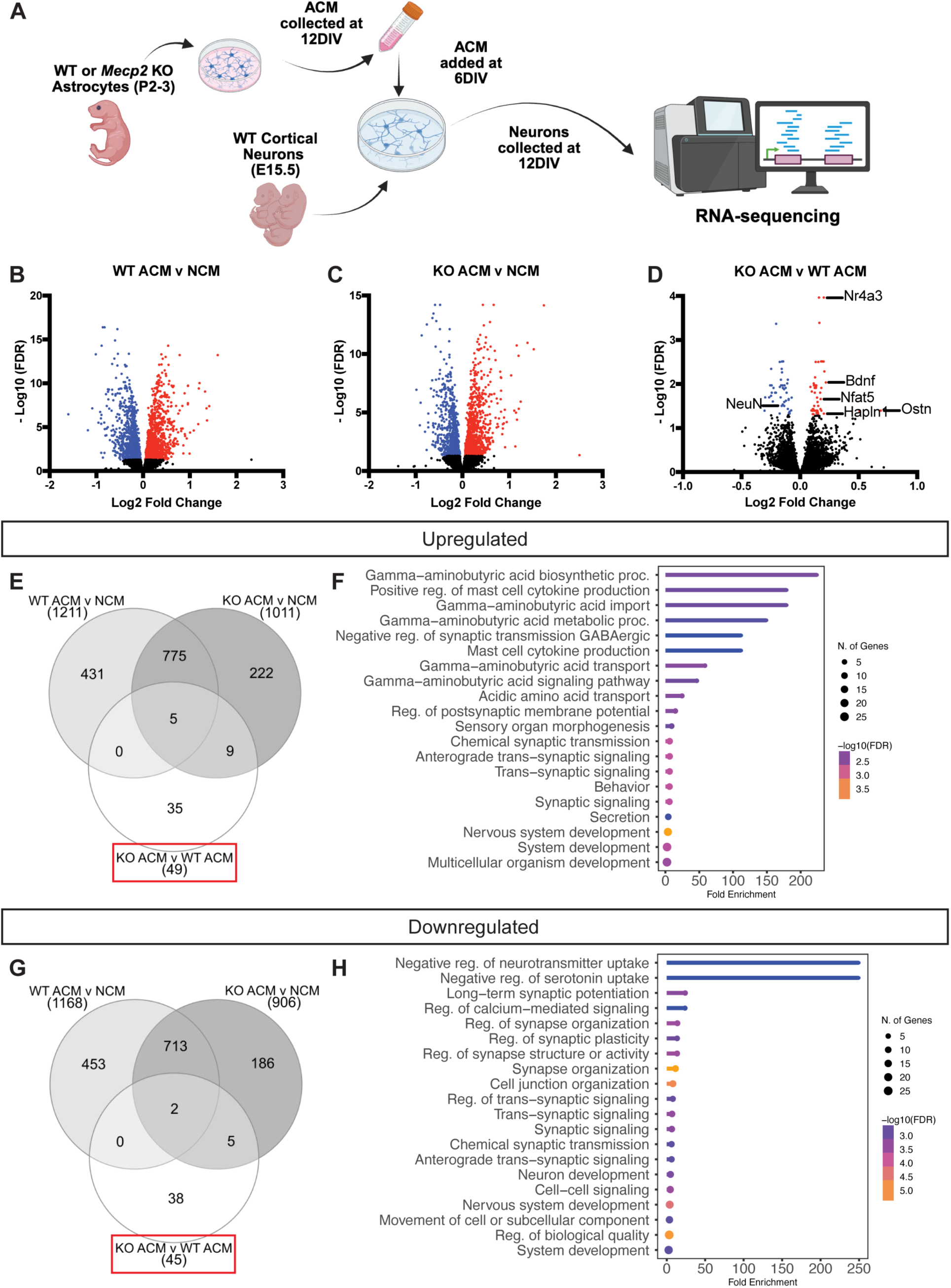
Astrocyte Conditioned Media (ACM) from *Mecp2* KO mice leads to transcriptome changes in cortical neuronal cultures compared to those treated with *Mecp2* WT ACM. (A) Schematic of experimental design. Wildtype CD1 E15.5 cortical neurons were treated with 20% v/v ACM from *Mecp2* WT or *Mecp2* KO astrocytes from 6-12 DIV, or with non-conditional media (NCM). RNA was collected from neurons on 12 DIV and bulk RNA-sequencing performed. (B-D) Volcano plots depicting differentially regulated genes (DEGs) between cortical neuronal cultures treated with (B) WT ACM compared to NCM, (C) KO ACM compared to NCM, or (D) KO ACM compared to WT ACM. KO ACM treatment leads to upregulation of 49 genes (red) and downregulation of 45 genes (blue) when compared to WT ACM (Genes of interest are denoted; FDR cut-off = 0.05). (E, G) Venn diagrams depicting numbers of DEGs between all three comparison groups: (E) upregulated DEGs or (G) downregulated DEGs (red boxes denote gene lists analyzed for gene ontology). (F, H) Gene Ontology analyses of Biological Processes enriched in DEGs between cortical neuronal cultures treated with *Mecp2* KO) ACM compared to *Mecp2* WT ACM: (F) upregulated DEGs and (H) downregulated DEGs. The top 20 most significantly upregulated categories ranked by FDR and fold enrichment are indicated. (Shiny Go 8.0; FDR cut-off 0.05)

We treated E15.5 CD1 wildtype cortical neurons with ACM collected from astrocytes cultured from P2-3 *Mecp2* WT or *Mecp2* KO cortex, in comparison to non-conditioned control media (NCM) from 6-12 DIV. We confirmed that *Mecp2* KO ACM leads to decreased neuronal outgrowth of wildtype neurons by 12 DIV (Supplemental Fig. 1A-C) as previously reported (Ballas et al., 2009; Williams et al., 2014; Caldwell et al., 2022; Albizzati et al., 2024). ACM from both *Mecp2* WT and *Mecp2* KO cultures induced extensive transcriptome changes in neurons in comparison to non-conditioned media (Fig. 3B-D), similar to what was observed in co-culture experiments (Albizzati et al., 2024). We focused, however, on the transcriptome changes observed with KO ACM in comparison to WT ACM (Fig. 3D). We identified 94 differentially expressed genes when comparing neurons treated with KO ACM versus WT ACM, 49 of which were up-regulated (Fig. 3E-F; Supplemental Table 1) and 45 of which were down-regulated (Fig. 3G-H; Supplemental Table 1). Gene Ontology analyses identified significant up-regulation of cytokine categories and GABAergic development (Fig. 3F; Supplemental Table 2), while transcript abundance of NeuN was down-regulated along with gene ontology categories related to long-term synaptic potentiation, neurotransmitter uptake, and synapse organization and plasticity (Fig. 3H; Supplemental Table 3).

We confirmed enhanced expression of GAD65/67 and reduced expression of NeuN with *Mecp2* KO ACM. Interestingly, protein expression levels of GAD65/67 and NeuN are not significantly different in *Mecp2* KO ACM-treated neurons at 12 DIV (Supplemental Fig. 1D-F), when transcript levels were found to be altered in the RNA-seq data; however, there is reduced expression of NeuN and enhanced expression of GAD65/67 by 16 DIV with *Mecp2* KO ACM (Supplemental Fig. 1G-I), perhaps reflecting a delay in the protein translation. These data, however, align with the RNA-seq findings and they suggest that the neurons treated with *Mecp2* KO ACM are altered in their developmental trajectory.

Importantly, *Mecp2* KO ACM leads to enhanced expression of the key PNN component *Hapln1* (hyaluronan and proteoglycan link protein 1) by wildtype neurons. PNN formation in the early postnatal brain initializes with the upregulation of HAPLN1 and synthesis of the HA backbone and mice deficient in HAPLN1 display severely attenuated PNN structure and formation (Carulli et al., 2010; Kwok et al., 2010). Exogenous over-expression of HAPLN1 accelerates maturation of CA1 PNNs and produces adult-like memory phenotypes (Ramsaran et al., 2023). These data suggest that *Mecp2* KO astrocytes might lead to enhanced expression of *Hapln1* by neurons, contributing to the precocious PNN formation.

### Mecp2 KO ACM causes enhanced HAPLN1 expression and precocious PNN formation on wildtype neurons

To investigate whether *Mecp2* KO ACM leads to enhanced HAPLN1 expression and PNN formation in wildtype neuronal cultures, primary astrocyte cultures were prepared from P3 WT and KO cortex with ACM added to wildtype E15.5 cortical neuronal cultures from 6-12 DIV, as performed for the RNA-seq experiments (Fig. 3A). The cultures were treated with AraC to eliminate glia from 1-3 DIV before treating with ACM from 6-12 DIV, after which expression of HAPLN1, and the additional PNN / ECM components ACAN, and TNR was compared between NCM (Fig. 4A-A’’), WT ACM (Fig. 4B-B’’), and KO ACM (Fig. 4C-C’’). Notably, the *Mecp2* WT ACM did not significantly alter the number of neurons displaying HAPLN1 (Fig. 4D), ACAN (Fig. 4E), or TNR positive PNNs (Fig. 4F) when compared to control non-conditioned media (NCM). This could suggest that secreted factors from WT astrocytes do not drive the change observed in PNN formation found with astrocyte co-culture (Fig. 2). *Mecp2* KO ACM, on the other hand, lead to a significant increase in PNN-expressing neurons, when compared to NCM and WT ACM, indicating that secreted factors from *Mecp2* KO astrocytes can enhance PNN formation, possibly through the regulation of *Hapln1* expression within the neurons. The number of HAPLN1 expressing neurons remains enhanced at 16 DIV with *Mecp2* KO ACM (Supplemental Fig. 2). Together, these data suggest that *Mecp2*-null astrocytes play a key role in the precocious formation of PNNs in RTT, providing essential insight into the mechanisms and structure of aberrant PNNs in *Mecp2*-null cortex and the early closure of critical period plasticity observed in RTT.

**Figure 4.**
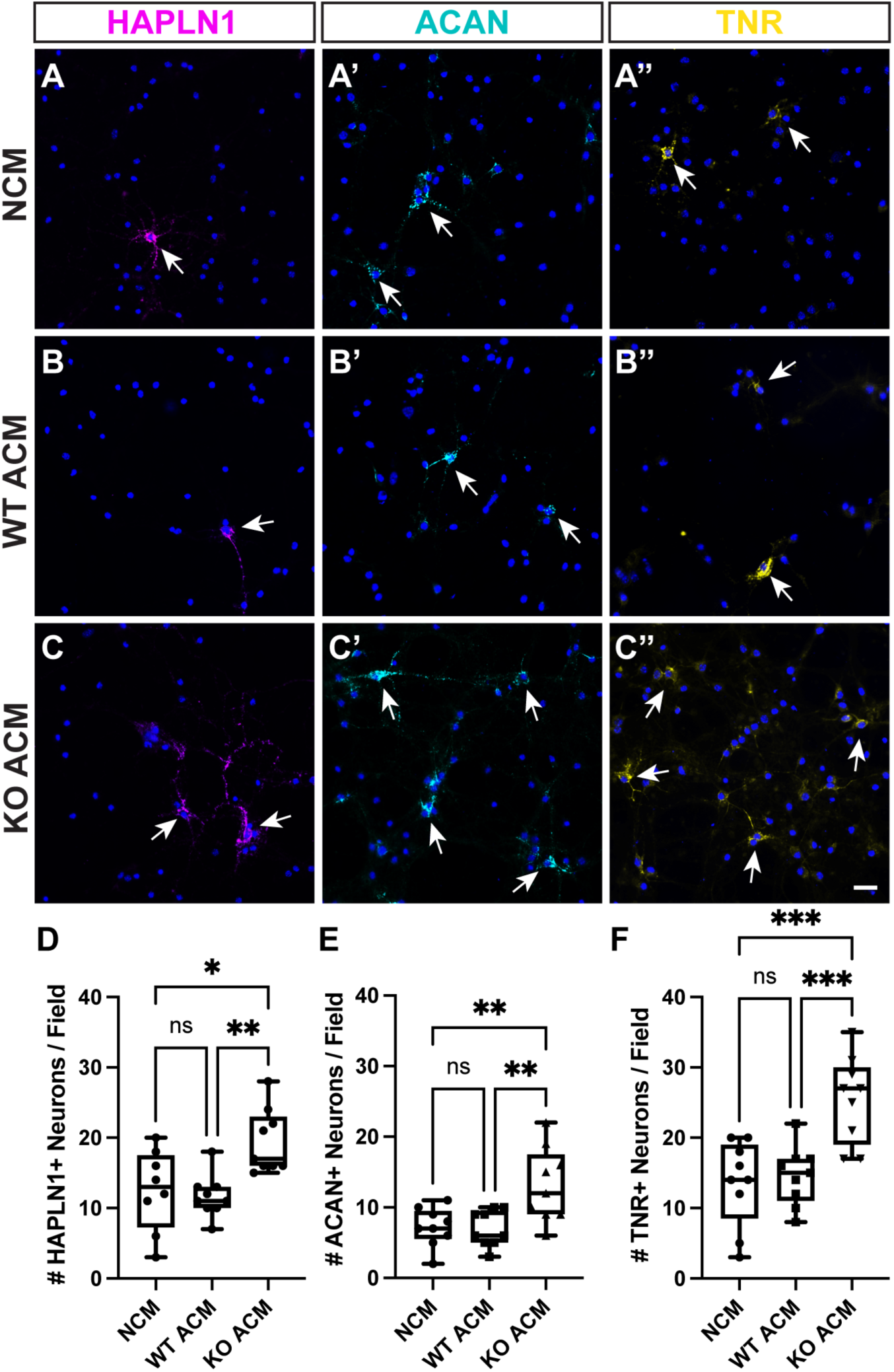
Enhanced formation of PNNs on wildtype cortical neurons treated with KO ACM. (A-C) ACM from WT (B) or KO (C) astrocyte culture media was collected at 12 DIV and was added to neuronal growth media at 20% v/v at DIV6 – DIV12 of E15.5 WT cortical cultures, in comparison to non-conditioned media control (A; NCM). Neuronal cultures were treated with AraC from days 1-3 to inhibit glia. Immunohistochemistry was performed for PNN / ECM components ACAN, HAPLN1, and TNR. PNN positive neurons are indicated with arrows. (D-F) The number of ACAN (D), HAPLN1 (E), and TNR (F) positive neurons was quantified from 3 large scan images per condition from each of 3 separate cultures. *Mecp2* WT ACM does not significantly alter the number of PNN-expressing neurons in comparison to NCM; however, *Mecp2* KO ACM significantly increases the number of PNN-expressing neurons relative to both NCM and WT ACM, suggesting enhanced PNN formation with KO ACM. * p<0.05, ** p<0.01, *** p<0.001. One-way ANOVA with Tukey’s post hoc test. Scale bars = 20μm. Error bars: mean ± S.D.

### HAPLN1 expression is increased in developing Mecp2 KO cortex

We next investigated whether HAPLN1 expression is also increased in the developing *Mecp2* KO neocortex *in vivo*. While precocious PNN formation and early critical period closure are well-established in RTT, to date studies in *Mecp2*-mutant mice have been limited to detection of PNNs by staining with Wisteria floribunda agglutinin (WFA), a lectin considered to be a standard reagent to detect PNNs, but that does not detect all PNNs. PNNs are composed of the glycosaminoglycan hyaluronan (HA), chondroitin sulfate proteoglycans (CSPGs), and secreted glycoproteins, but an understanding of their precise composition is incomplete (for review see (Lau et al., 2013)). We determined that HAPLN1 expression is significantly increased in the developing *Mecp2* KO neocortex at P16, relative to WT littermates (Fig. 5A-A’). We found that Brevican (BCAN), another key component of the neural ECM, and PNNs specifically, is also significantly increased while TenascinR (TNR) is not. Importantly, there are no significant differences in expression of the three PNN / ECM components at P21 (Fig. 5B-B’) suggesting that there is a developmental shift in PNN component expression rather than a sustained increase in expression.

**Figure 5.**
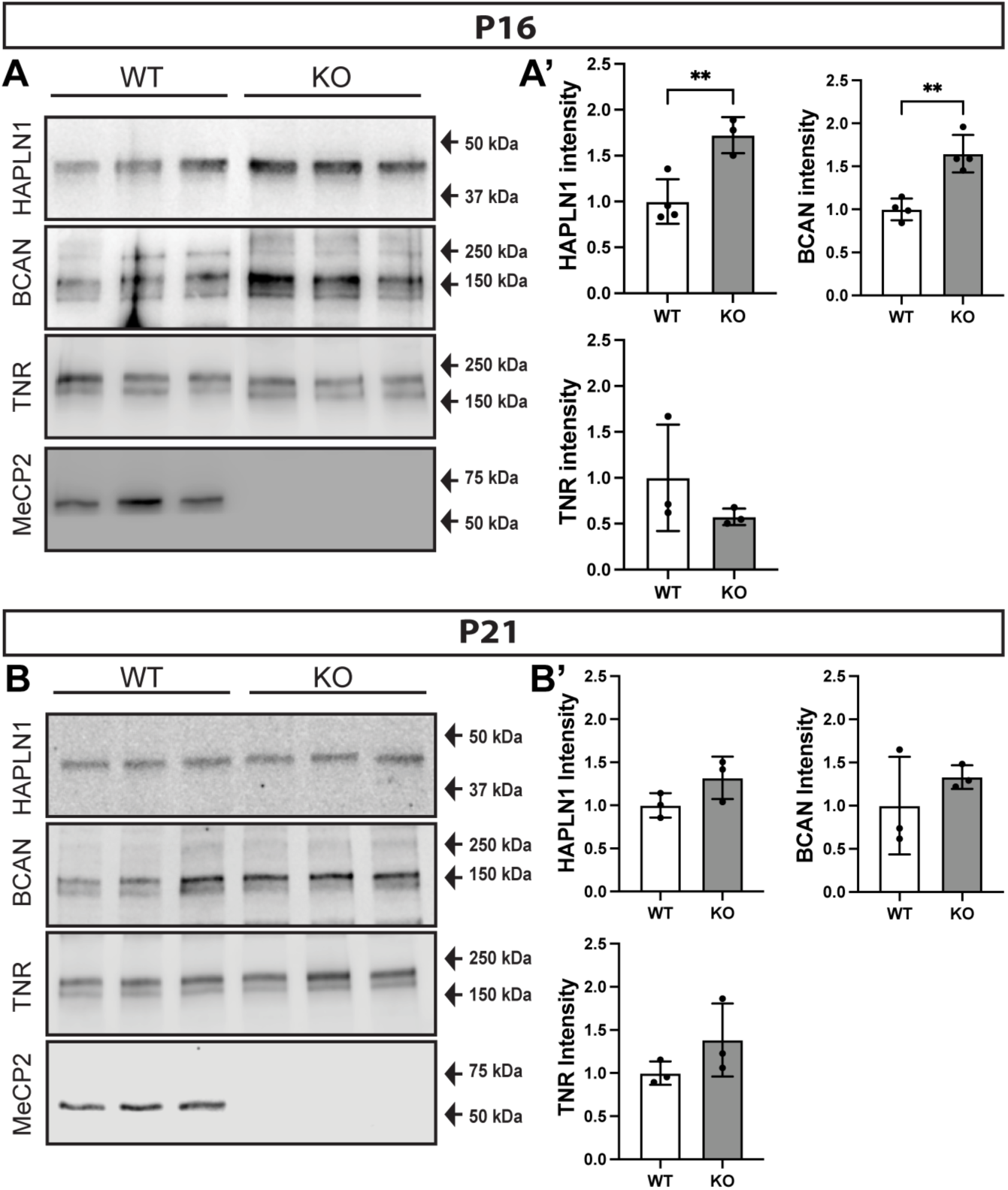
PNN / ECM components are upregulated in *Mecp2* KO cortex compared to *Mecp2* WT at P16. (A,B) Western blot analysis of (A) P16 and (B) P21 cortical homogenates. (A) Hapln1 and Brevican are upregulated in KO brains compared to WT brains at P16, whereas TenascinR (TNR) shows no change. (B) PNN / ECM components show no statistically significant upregulation at P21. (A’,B’) Fluorescent intensities normalized to total protein and then compared to WT levels for n = 3-4 animals per developmental stage per genotype; two-tailed T-test, ** p < 0.01.

### Altered Developmental Trajectory of PNN Structure in the Mecp2 KO neocortex

Neural ECM, including the highly organized PNN structure, undergoes dynamic changes during development. PNN components are immobilized on the surface of the cell by two independent types of interactions; PNN formation initializes with the synthesis of a hyaluronic acid (HA) backbone and HAPLN1 expression (Carulli et al., 2010), followed by a Ca^2+^ dependent incorporation of other PNN components, including ACAN, TNR, and RPTPζ (Carulli et al., 2010; Eill et al., 2020; Sinha et al., 2023). Although TNR expression is not altered in the P16 KO cortex, as HAPLN1 and Brevican are, TNR is a key ECM organizer known to promote the assembly and incorporation of ECM molecules via the clustering of CSPGs (Bruckner et al., 2000; Haunso et al., 2000; Morawski et al., 2014; Suttkus et al., 2014). TNR associates with the cell membrane by its interaction with HA via its clustering of CSPGs and through Ca^2+^ mediated protein-protein interactions. We thus sought to determine if the developmental trajectory of TNR incorporation is disrupted in the *Mecp2* KO neocortex.

To first establish the developmental timing of incorporation of TNR in the neocortex, we isolated membrane fractions from P21, P36 and P56 cortex homogenates. We disrupted the HA backbone using the enzyme chondroitinase (ChABC) and determined the ratio of TNR in the solubilized (released, R fraction) fraction relative to the pelleted fraction (i.e. proteins still attached to the membrane, P fraction) by Western blots (Fig. 6A). As a control, we examined HAPLN1, an ECM protein known to associate with the cell surface predominantly via its interaction with HA (Fig. 6B). These were then compared to the release that occurred with the combination of ChABC and EDTA to additionally disrupt the Ca^2+^ dependent protein-protein interactions. The release of TNR with ChABC treatment from wildtype cortex shows a gradual decrease in solubility from P21 to P56, indicative of its incorporation into the ECM along with ECM maturation (Fig. 6A,A’); the release of TNR at P21 is significantly greater than that at P56 (Fig. 6A’). This decreasing trend in release of TNR, however, is not observed when ChABC treatment is combined with EDTA indicating that TNR increasingly associates with the cell membrane in a Ca^2+^ dependent mechanism during development (Fig. 6C-C’). HAPLN1 release is comparable between treatments with ChABC and ChABC along with EDTA (Fig. 6B’), which is consistent with the model that HAPLN1 is incorporated into the ECM in a predominantly HA-dependent manner (Fig. 6C-C’).

**Figure 6.**
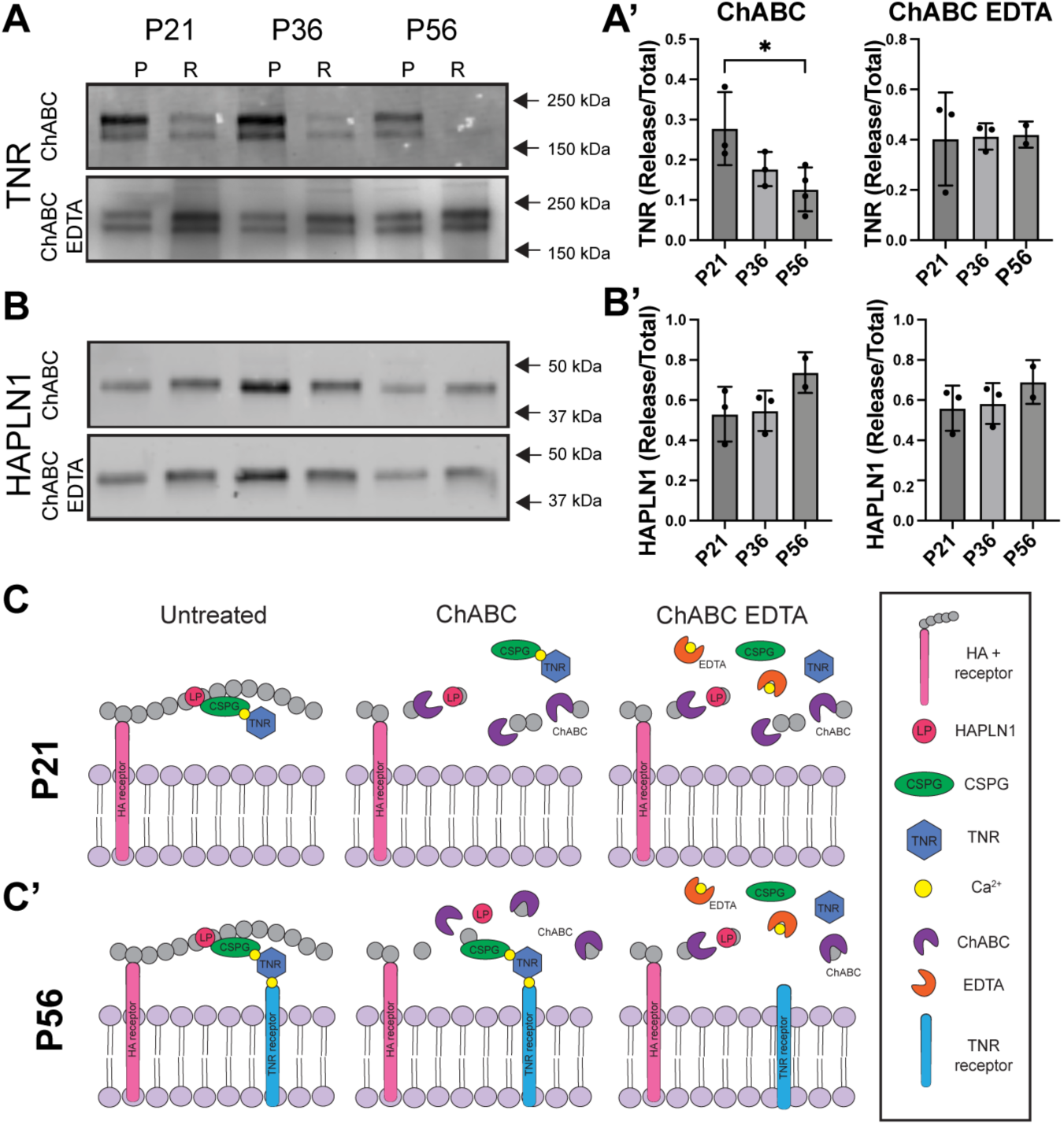
PNN / ECM component TNR binds differentially to the cell surface across development. (A-B) Membrane fractions were isolated from P21, P36 and P56 cortex homogenates. We disrupted the HA backbone using the enzyme chondroitinase (ChABC) and determined the ratio of TNR in the solubilized (released, R fraction) fraction relative to the pelleted fraction (i.e. proteins still attached to the membrane, P fraction) by Western blots. These were then compared to the release that occurred with the combination of ChABC and EDTA to additionally disrupt the Ca2+ dependent protein-protein interactions. (A,A’) There is modest release of TNR (27 ± 09%) into the soluble fraction (R) at P21 with ChABC treatment alone. Release of TNR is lower at P36 (17 ± 04%) and significantly lower at P56 (12 ± 05%; * p = 0.041, Ordinary one-way ANOVA, Tukey’s multiple comparisons test). The majority of TNR remains behind in the insoluble pellet (P) at this stage. This effect is not seen when ChABC treatment is combined with EDTA. (B,B’) HAPLN1 associates with the cell membrane primarily in a HA dependent manner and is not differentially released into the soluble R fraction by ChABC or ChABC and EDTA treatment. n = 3-4 animals per developmental stage. (C,C’) We propose a model of the ECM in which during early postnatal development, HAPLN1, CSPGs and TNR are linked to the cell surface in a HA dependent manner (C). As the animal matures (C’), TNR further associates cell surface receptors in a Ca2+ dependent mechanism. Accordingly, ChABC treatment to digest HA releases TNR from membranes during early postnatal development (P21) and not during later stages (P56). A combination of ChABC treatment and EDTA to chelate Ca2+ ions releases TNR into soluble phase at PND56.

We next compared the release of TNR and HAPLN1 between *Mecp2* WT and KO cortex at P21 (Fig. 7A-B), when TNR demonstrates incomplete developmental Ca^2+^ dependent incorporation in the WT. We found that the release of TNR with ChABC treatment from *Mecp2* KO cortex was significantly lower than that of WT cortex (Fig. 7A-A’), although this this difference was eliminated when ChABC treatment was combined with EDTA as noted in the WT. We confirmed that no difference in release was observed for HAPLN1 between the two genotypes with ChABC or ChABC+ EDTA treatment (Fig. 7B-B’). At P36, no significant differences in release of TNR (Fig. 7C-C’) or HAPLN1 (Fig. 7D-D’) were observed. These findings indicate that the Ca^2+^ dependent incorporation of TNR is significantly higher in the P21 *Mecp2* KO cortex, suggesting a structural maturation similar to a mature P36 or P56 WT TNR incorporation (Fig. 7E). Together, these studies suggest a precocious formation and maturation of PNNs in the *Mecp2* KO neocortex.

**Figure 7.**
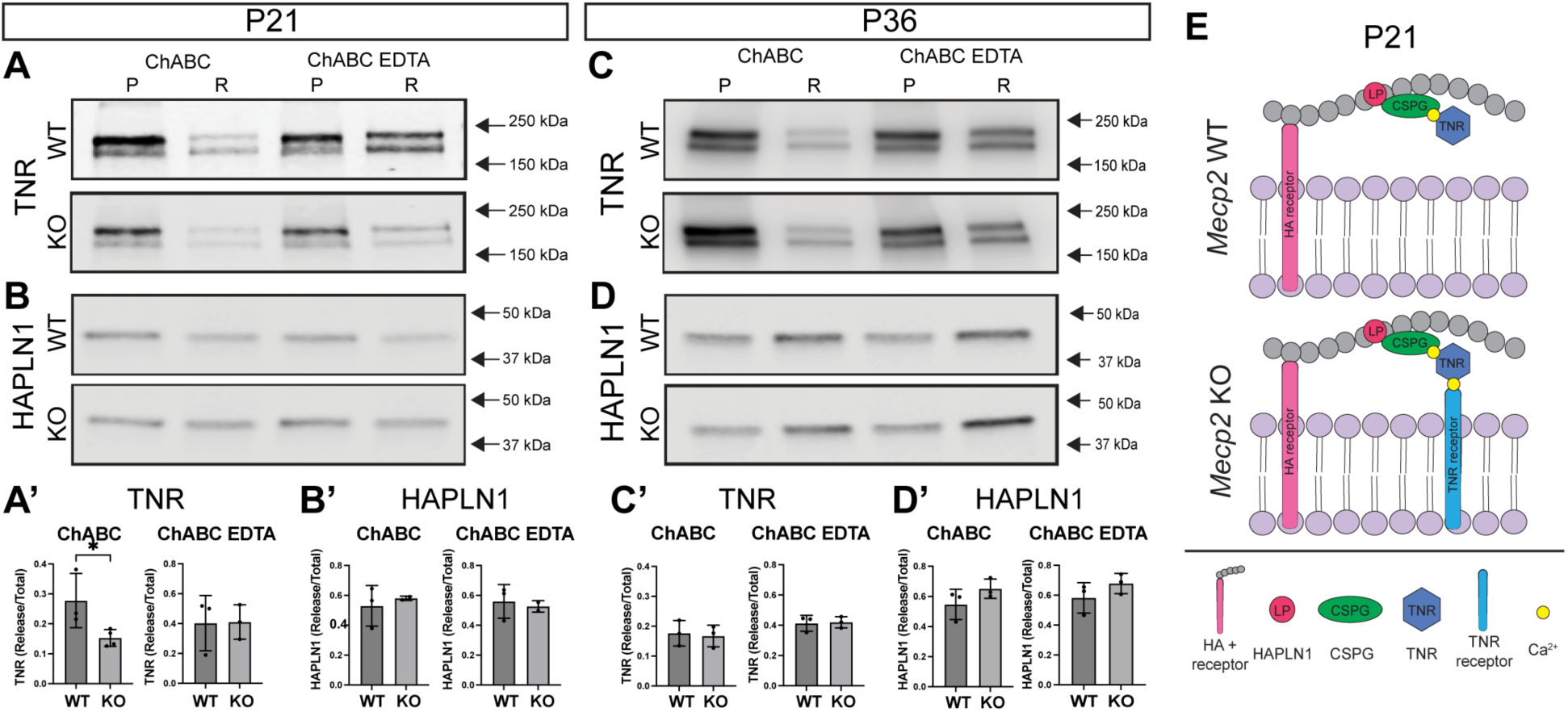
Ca2+ dependent incorporation of TNR to the cell membrane is greater in *Mecp2* KO cortex at P21. (A-B) Western blots and quantification showing release of TNR (A) and control molecule HAPLN1 (B) from isolated membranes from WT and KO mouse cortex at P21. (A’) Compared to WT animals (27 ± 09%), release of TNR into the soluble fraction (R) is significantly lower in KO cortex (15 ± 02%; p = 0.044) after ChABC treatment, but not when ChABC treatment is combined with EDTA. (B’) The differential release after ChABC or ChABC and EDTA treatment is not observed for HAPLN1. (C-D) Unlike at P21, release of TNR into the soluble (R) fraction is not significantly different between WT and KO mouse cortices with ChABC treatment (C’). (D’) HAPLN1 is not differentially released with ChABC or ChABC EDTA treatment at P36, as also observed at P21. N = 3-4 animals per developmental stage per genotype. (E) Schematic showing association of TNR with the cell surface in *Mecp2* WT and *Mecp2* KO brains at P21. TNR incorporates more tightly with the cell membrane in a Ca2+ dependent mechanism in KO at this early developmental stage, with a similar profile to that observed at P56 in the WT.

## Materials and Methods

### Animals

All animal experimental protocols were approved by the Syracuse University Institutional Animal Care and Use Committee and adhere to NIH ARRIVE guidelines. Mice were group housed at a maximum of five mice per cage on a 12/12 h light/dark cycle and were given food and water ad libitum. CD-1 timed pregnant female mice were purchased from Charles River. Female *Mecp2* heterozygous mice were purchased from The Jackson Laboratory (B6.129P2(C)-Mecp2tm1.1Bird/J; RRID:IMSR_JAX:003890) and were maintained on a C57BL/6 background. Genotypes were determined by PCR on genomic DNA as follows: *Mecp2* mutant mice, forward primer oIMR1436 5’ - GGT AAA GAC CCA TGT GAC CC - 3’; reverse primer oIMR1437 5’ - TCC ACC TAG CCT GCC TGT AC - 3’; reverse primer oIMR1438 5’ - GGC TTGCCACATGACAA - 3’.

### Primary cortical neuronal cultures

Embryonic day (E)15.5 embryos were collected from timed pregnant CD-1 mice (Charles River, 022) and the cortices were dissected out in dissociation medium (DM) containing MgKyn, glucose, AP-V (Sigma-Aldrich, A5282), penicillin-streptomycin (Invitrogen, 15070063), and B27 supplement (Gibco, 17504044). Harvested tissue was dissociated using cysteine (Sigma-Aldrich, C9768) and papain (Sigma-Aldrich, P3125) in OptiMem media (Gibco, 51985034). Dissociated cells were plated on glass coverslips precoated with poly-D-lysine hydrobromide (Sigma-Aldrich, P0899) at a density of 750,000 cells/mL. Cells were initially plated in Neurobasal based medium containing Glutamax (Gibco, 21103049), fetal bovine serum (Invitrogen, 50253931), and penicillin-streptomycin (Gibco, 15140122). After 4 hours, the plating medium was removed, and growth medium containing Neurobasal (Gibco, 21103049), Glutamax (Gibco, 35050061), penicillin-streptomycin (Invitrogen, 15070063), N2 (Gibco, 17502048) and B27 supplements (Gibco, 17504044) was added. Cells were treated with AraC (Sigma-Aldrich, C6645) from DIV 1 to DIV 3 to remove glia as required. A full media change was carried out on DIV 3 to remove AraC and subsequently, half media changes were carried out every 3 days. ACM treatment was started at 6 DIV with addition of 20% ACM by volume to individual wells.

### Primary cortical astrocyte cultures

Cortical astrocyte cultures were prepared from P2-3 mice brains as described previously (Uliasz et al., 2012). Briefly, P2 cortices were dissected out in cold dissociation media and meninges were removed. Tissue was minced and digested using 0.025% Trypsin EDTA for 20 minutes. Cells were triturated using glass pasteur pipettes and plated on Primaria plates. A full media change was carried out 24 hours after plating. Subsequently, cells were given half media changes every 3 days. Conditioned media from cells (ACM) were collected at 12DIV and stored at –80°C freezer for future use.

### Immunocytochemistry

Coverslips were fixed in cold 4% phosphate-buffered paraformaldehyde (PFA) with 0.01% glutaraldehyde, pH 7.4. Cells were then blocked in screening medium (DMEM, 20% FBS, 0.2% sodium azide) for 1 hour, before adding primary antibodies overnight at 4°C. The following day, Alexa-fluor conjugated secondary antibodies in screening medium were added to the cells for 2 hours before mounting the coverslips using Fluoromount (Southern Biotech) containing DAPI for visualizing cell nuclei. Primary antibodies used were mouse anti-NeuN (Millepore, MAB377), rabbit anti-Gad1/Gad2 (Sigma, G5163), mouse anti-TenascinR (R&D Sytems, MAB1624), goat anti-HAPLN1 (R&D Systems, AF2608), and rabbit anti-ACAN (Millepore, AB1031).

### Quantification

The number of cells positive for each PNN marker was counted from large scan images (1769μm X 1769μm) from 3 different areas for each coverslip using a Nikon Eclipse Ni upright microscope with a Prime BSI Express sCMOS camera (Teledyne Photometrics) under a 20X objective, equipped with an OptiScan XYZ motorized stage (Prior Scientific) to allow for Z stacks, and NIS Elements (Nikon) software. Counts were performed blinded to condition, and verified by an independent investigator.

### Immunohistochemistry

Brains were perfused, fixed in 4% paraformaldehyde overnight, washed in PBS, and then cryopreserved with increasing concentrations of sucrose in PBS (5%, 15%, 30%) until the brains settled. Brains were then frozen in Tissue-Tek O.C.T. Compound (Avantor, 25608930). The cryoblocks were cryosectioned for 30 µm thick coronal sections, with brain sections then being stored in PBS at 4°C until immunolabeling. As a broad PNN marker, Fluorescein labeled WFA (Wisteria floribunda agglutinin) from Vector Laboratories Inc. (FL-1351-2) was used.

### Biochemical release assay

The neocortex was dissected and tissue was homogenized in 150mM sodium chloride and 50mM Tris with EDTA-free protease inhibitor tablets (Roche, Indianapolis, IN, USA 1 tablet in 10mL buffer), using a handheld homogenizer. Homogenates were centrifuged at 8,000g for 10 min at 4 °C. The supernatant was then removed, the pellet washed once and then resuspended in 1 mL buffer. A Bradford (Bio-Rad) assay was performed, and protein concentrations were adjusted to 2.0 μg/μL. Samples were treated with 2 µL chondroitinase ABC (0.01 U/ µL, Sigma-Aldrich, Saint Louis, MO, USA) per 100 µl of sample and/or 2.5 mM EDTA for 8 hours. Samples were centrifuged again at 8,000g for 10 min at 4 °C to separate soluble release (R) fraction and insoluble pellet (P) fractions and subsequently prepared for Western blotting by adding sample loading buffer and heating to 95 °C for 5 minutes.

### Western Blotting

Protein concentrations were determined by Bradford assay before gel electrophoresis. 6-15% gradient SDS-polyacrylamide gels were used and transferred to 0.45 μM nitrocellulose membranes. Western blotting was conducted as previously described (Eill et al., 2020; Sinha et al., 2025). Briefly, blots were blocked in 5% milk in low salt TBST and then incubated in primary antibody overnight. Primary antibodies used were rabbit anti-MeCP2 (Cell Signaling, 3456), mouse anti-TenascinR (R&D Sytems, MAB1624), goat anti-HAPLN1 (R&D Systems, AF2608), and rabbit anti-Brevican (B756, provided by Dr. R.T. Matthews). Blots were then incubated in HRP-conjugated secondary antibodies (Jackson Laboratory, Bar Harbor, ME, USA) and exposed using SuperSignal west pico chemiluminescent substrate (Thermo-Fisher Scientific, Rockford, IL, USA) or incubated with IRDye 800CW and directly imaged (LI-COR Biotech, Lincoln, NE, USA). Blots were imaged using the LI-COR Odyssey Fc imaging system (LI-COR Biotech, Lincoln, NE, USA). Loading concentrations were judged by total protein using REVERT Total Protein Stain 700 (LI-COR Biotech, Lincoln, NE, USA).

### RNA-sequencing

Total RNA was extracted from 12DIV neuronal cultures treated with non-conditioned media (NCM), *Mecp2* WT ACM, and *Mecp2* KO ACM using a combined TRIzol reagent (Invitrogen) and RNeasy Mini Kit (Qiagen) protocol, as described previously (Ribeiro and MacDonald, 2022). Samples were from 4 independent neuronal cultures. RNA samples were assessed for quality and concentration by NanoDrop (ThermoFisher Scientific) and Agilent Tapestation (Agilent Technologies). Library preparation and RNA-sequencing were performed by Azenta Life Sciences. Libraries were generated with the NEBNext Ultra RNA Library Prep Kit for Illumina following manufacturer’s instructions (New England Biolabs), and sequenced on an Illumina Hi-seq Series (2x150bp) with 20 million paired-end reads per sample. Sequence reads were trimmed to remove adapter sequences using Trimmomatic v.0.36 and were aligned to the Mus Musculus GRCm38 ERCC reference genome using the STAR aligner v.2.5.2b. Gene hit counts were extracted using featureCounts from the Subread package v.1.5.2. Sequence reads underwent batch correction to remove unwanted variation in R using RUVseq (Risso et al., 2014) and differentially expressed genes were calculated using edgeR (Robinson et al., 2010). Genes were considered differentially expressed according to their false discovery rate (FDR < 0.05). Volcano plots were made using GraphPad Prism and gene ontology (GO) performed using ShinyGo 0.8 (Ge et al., 2020).

### Statistical analysis

GraphPad Prism 10 (GraphPad Software) was used to carry out the statistical analyses. No statistical method was used to predetermine sample sizes, but our sample sizes are similar to those generally employed in the field. Our statistical tests consisted of two-tailed t test, or two-way ANOVA or mixed-effects analysis with Tukey’s multiple comparison, or Pearson correlation of multivariate analysis to determine statistical significance among groups. Data distribution was handled as if normal, but this was not formally tested (since potential differences in results would be minor). All data shown represent mean ± SD. Sample size and statistical test are specified in each figure legend. Final images were assembled into figures using Adobe Photoshop and Adobe Illustrator.

## Discussion

Our findings identify a non-cell-autonomous role for astrocytes in the precocious formation of PNNs and structural maturation of ECM during postnatal development in the *Mecp2* KO cortex. PNNs form the most intricate and conspicuous extracellular macromolecular structures in the CNS. They are essential in the modulation of neuronal plasticity and brain maturation (Wang and Fawcett, 2012). PNNs form precociously and the critical period of plasticity closes prematurely in RTT, likely limiting the developmental refinement of circuitry. The *Mecp2* KO visual cortex displays accelerated maturation of parvalbumin (PV) interneurons (Patrizi et al., 2020), accelerated onset and closure of the critical period for ocular dominance plasticity (Krishnan et al., 2015) and precocious appearance of PNNs (Krishnan et al., 2015; Sigal et al., 2019; Patrizi et al., 2020). Similarly, PNNs form precociously and remain enhanced in CA2 of the hippocampus of *Mecp2* KO mice, prematurely restricting LTP plasticity (Carstens et al., 2021). PNNs are also aberrantly increased in auditory and somatosensory cortex of female heterozygous (*Mecp2*+/-) mice in response to learned maternal behavior (Krishnan et al., 2017; Lau et al., 2020). Preventing PNN enhancement during this learned behavior improves the deficient pup retrieval behavior in *Mecp2*+/-mice, suggesting that the increased PNNs inhibit auditory plasticity that is necessary for this learned behavior (Krishnan et al., 2017).

The enzymatic disruption of PNNs can partially restore plasticity in adults and improve memory (Bruckner et al., 1998; Pizzorusso et al., 2002; Romberg et al., 2013; Yang et al., 2015), highlighting the exciting potential to modulate PNN development and structure to prevent the precocious closure of the critical period plasticity, or to reopen it, toward improvement of RTT phenotypes. PNNs are composed of the glycosaminoglycan hyaluronan (HA), chondroitin sulfate proteoglycans (CSPGs), and secreted glycoproteins, but an understanding of their precise composition is incomplete (for review see (Lau et al., 2013)). To date, studies in *Mecp2*-mutant mice have been limited to detection of PNNs by staining with Wisteria floribunda agglutinin (WFA), a lectin considered to be a standard reagent to detect PNNs, but that does not detect all PNNs. WFA most likely binds to a chondroitin sulfate epitope dependent on the CSPG aggrecan, however, the precise mechanism of its binding to PNNs is unclear. Further, current studies have employed injections of chondroitinase ABC to digest the HA backbone of PNNs and reopen the critical period; this is a sledgehammer approach that disrupts the ECM broadly. There is little understanding of the molecular composition and structure of the aberrant PNNs formed in RTT, which is essential to develop strategies to specifically rescue their formation and elucidate mechanisms by which they disrupt plasticity.

Our data demonstrate precocious Ca^2+^-dependent incorporation of TNR into the ECM by P21 in the *Mecp2* KO cortex (Figs 6,7). TNR promotes the assembly of the ECM in the CNS, including PNNs, via clustering of CSPGs, and TNR null animals also show disrupted PNN structure (Bruckner et al., 2000, Haunso et al., 2000, Suttkus et al., 2014, Morawski et al., 2014). These disruptions, however, are distinct from mice lacking HAPLN1 (Haunso et al., 2000; Carulli et al., 2010; Kwok et al., 2010; Morawski et al., 2014; Eill et al., 2020), which shows enhanced expression in the P16 *Mecp2*-null cortex (Fig. 5). HAPLN1 plays a pivotal role stabilizing the HA backbone and cross-linking lectican family members, and exogenous over-expression of HAPLN1 accelerates maturation of CA1 PNNs and produces adult-like memory phenotypes (Ramsaran et al., 2023). Recent studies employing tagged HAPLN1 demonstrate more widespread detection of ECM and PNN structures around many neurons, including excitatory neurons (Lemieux et al., 2023; Sterin et al., 2024), suggesting more extensive roles for HAPLN1 in the neural ECM and brain maturation. Modulation of HAPLN1 expression may, therefore, offer a potential therapeutic target for more focused prevention or reversal of the aberrant PNN formation in RTT. Importantly, we identified increased expression of HAPLN1 by wildtype neurons in response to *Mecp2* KO ACM treatment (Fig. 3), indicating that modulation of HAPLN1 expression will likely require targeting of *Mecp2* KO astrocytes.

Interestingly, HAPLN1 promotes pro-inflammatory factors including IL-6 and NF-κB (Chen et al., 2022). Our RNA-seq data identified up-regulation of inflammatory cytokine production among the top Gene Ontology categories up-regulated by wildtype neurons in response to *Mecp2* KO ACM, in comparison to *Mecp2* WT ACM (Fig. 3). Further, a recent study indicates that IL-6 is the primary astrocytic factor that leads to disrupted synapse development in neurons cultured with *Mecp2* KO astrocytes (Albizzati et al., 2024). Our prior studies identified aberrant NF-κB signaling in *Mecp2*-mutant cortex and demonstrated improved neuronal complexity phenotypes of *Mecp2*-null neurons with NF-κB inhibition, both *in vivo* and *in vitro* (Kishi et al., 2016; Ribeiro et al., 2020). As neuroinflammation and oxidative stress disrupt PNNs (Cardis et al., 2018; Dwir et al., 2020; Phensy et al., 2020), these data suggest that reducing NF-κB and IL-6 signaling could also ameliorate PNN phenotypes. Further, they highlight astrocyte-neuron crosstalk as a key target.

Secreted factors from *Mecp2* KO astrocytes significantly disrupt neuronal development (Ballas et al., 2009; Williams et al., 2014; Caldwell et al., 2022) including neuronal complexity and PNN/ECM formation (Fig. 4). Interestingly, we found in a co-culture that wildtype astrocytes slowed the developmental trajectory of PNN formation on neurons, in comparison to cultures where astrocytes were largely eliminated (Fig. 2). However, *Mecp2* WT ACM did not significantly alter PNN/ECM formation on neurons when compared to control media, while *Mecp2* KO ACM lead to enhanced PNN/ECM formation on wildtype neurons by 12 DIV. These data suggest that secreted factors from *Mecp2* KO astrocytes specifically enhance PNN / ECM formation. The differential impact of co-culture with astrocytes versus secreted factors in ACM can also be observed in the recent study by Albizzati et al., where they found that enhanced secretion of IL-6 from *Mecp2* KO astrocytes only occurred with co-culture with neurons (Albizzati et al., 2024). Thus, enhanced PNN / ECM formation on neurons is likely the result of a complex interplay between astrocytes and neurons.

Previous studies have also identified a cell-autonomous role for the PV interneurons themselves in the formation and/or maintenance of PNNs. A PV-specific knockout of *Mecp2* using the Pvalb-Cre led to enhanced WFA-positive PNNs in the adult visual cortex (Patrizi et al., 2020). It should be noted that full removal of *Mecp2* from PV interneurons is not complete until P90 (Durand et al., 2012) and PNNs were analyzed at P200, so it is not clear if *Mecp2* is necessary in PV interneurons for the precocious developmental formation of PNNs or plays a role in their enhancement or maintenance (Patrizi et al., 2020). Using the same PV-Cre line, Rupert et al (Rupert et al., 2023) demonstrated that loss of *Mecp2* in PV interneurons was sufficient to recapitulate the enhanced PNN formation in the adult female Het auditory cortex in response to a learned pup retrieval task. These studies indicate that *Mecp2* expression in PV interneurons cell-autonomously contributes to PNN formation and/or maintenance in the adult brain. It will be interesting to determine if *Mecp2* expression in PV interneurons is necessary to prevent precocious formation and, conversely, whether *Mecp2* expression in astrocytes plays a role in the adult formation / maintenance of PNNs.

It should be noted that the current studies focused on male *Mecp2* KO mice to gain an understanding of how MeCP2 regulates the formation of PNNs. It will be important, however, to extend these studies to the more clinically-relevant female heterozygous *Mecp2* mice to determine whether they also demonstrate precocious maturation of the ECM. Further, such studies will enhance our understanding of whether the changes are cell-autonomous in the context of the cellular mosaicism of the heterozygous brain, where approximately half the cells express the wildtype allele and half the mutant allele due to random X-inactivation. Taken together, our data demonstrate that *Mecp2*-null astrocytes play a key role in the precocious formation of PNNs in RTT. These results provide essential insight into the mechanisms and structure of aberrant PNNs in *Mecp2*-null cortex and identify potential new avenues for targeted rescue or reversal of the precocious closing of the critical period in RTT.

## Acknowledgements

The authors thank Dr. Yasir Ahmed-Braimah (Syracuse University) for assistance with RNA-seq data analyses. Figure 3A was created with BioRender.com. This work was supported by the International Rett Syndrome Foundation [Grant #4021] and the National Institutes of Health [1R01NS106285] awarded to JLM.

## Author Contributions

AS, AMK, RTM, and JLM designed experiments; AS, AMK, and NK performed experiments; AS, AMK, NK, RTM, and JLM analyzed data; JLM, AS, and AMK wrote the manuscript with contributions by all authors.

