## Supplemental Materials for "Astrocytic Regulation of aberrant perineuronal net formation in *Mecp2*-null Neocortex"

### Supplemental Material

Supplemental Figures: 2

Supplemental Tables: 3

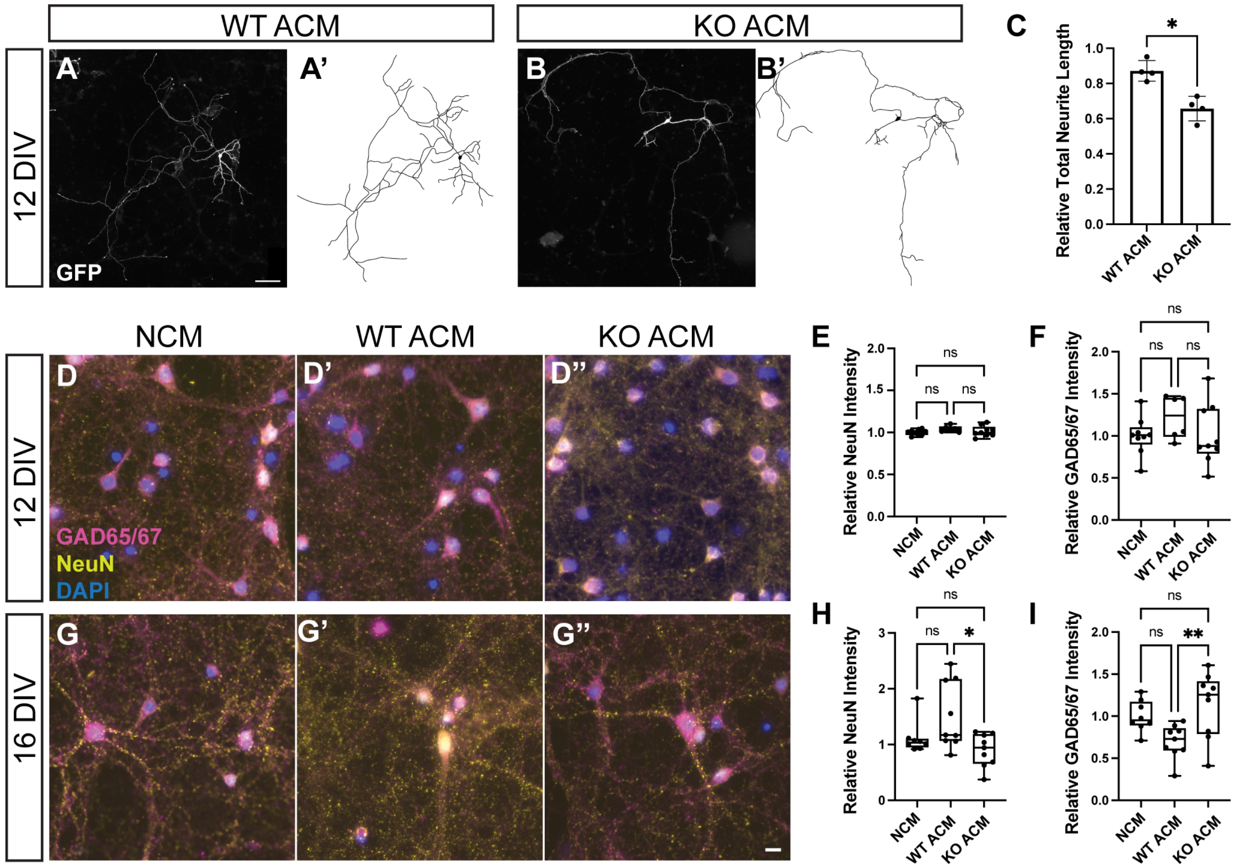


**Supplemental Figure 1. Astrocyte Conditioned Media (ACM) from *Mecp2* KO mice leads to changes in neuronal morphology and expression of Gad65/67 and NeuN.** E15.5 wildtype neuronal cultures were treated with ACM as outlined in Figure 3. (A-C) E15.5 wildtype neurons treated with *Mecp2* KO ACM show reduced total neurite length compared to *Mecp2* WT ACM. (A-B) Representative images and traces of wildtype neurons exposed to WT ACM (A-A') and KO ACM (B-B'). Scale bar = 50 μm. A minimum of 5 neurons from each treatment were traced and total neurite length measured from each of 4 independent cultures. Relative neurite length, normalized to the NCM control, was averaged for all neurons from each of the 4 independent cultures, showing a significant reduction with KO ACM (N = 4 cultures, two-tailed T-test, ** $p < 0.05$). (B-I) RNA-sequencing showed an upregulation of GABA related genes such as Gad67 and a downregulation of a mature neuronal marker, NeuN, with KO ACM at 12 DIV. (B-F) A significant change is not observed in either with immunohistochemistry at 12 DIV. (G-I) However, at 16 DIV, NeuN protein expression is downregulated in wildtype neurons treated with KO ACM, and GAD65/67 expression is upregulated. (Relative Intensity normalized to non-conditioned media (NCM) across 3 large scans (1769 μm x 1769 μm) per culture per developmental stage per condition (3 independent cultures); Scale bar = 10 μm, one-way ANOVA, Tukey's multiple comparisons test.

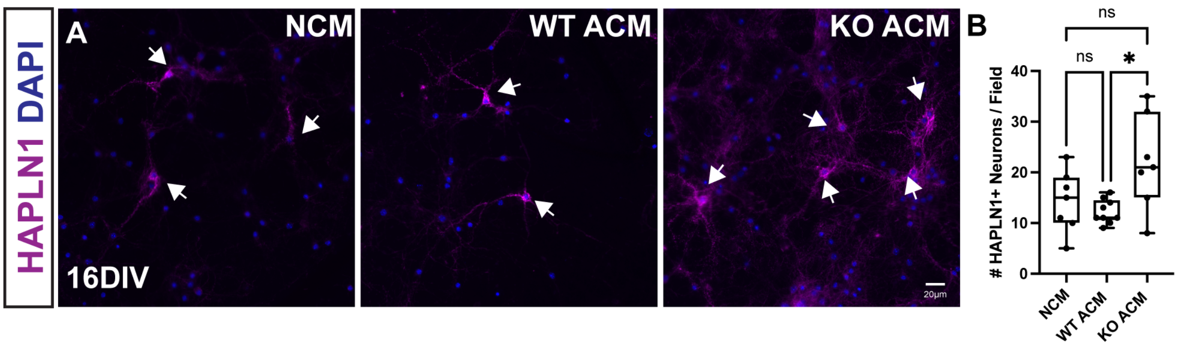


**Supplemental Figure 2: Upregulation of HAPLN1 with addition of Astrocyte Conditioned Media (ACM) from *Mecp2* KO mice persists to 16 DIV.** (A) E15.5 wildtype neurons were treated with *Mecp2* WT or *Mecp2* KO ACM from 6-16 DIV. Arrows denote HAPLN1+ neurons. (B) At 16 DIV, wildtype neurons treated with *Mecp2* KO ACM displayed an increased number of HAPLN1+ neurons compared to neurons treated with *Mecp2* WT ACM and non-conditioned media (NCM). * $p<0.05$. One-way ANOVA with Tukey's post hoc test. Scale bar = 20μm. Error bars: mean ± S.D.

**Supplemental Table 1:** Significant DEGs (FDR<0.05) in wildtypeneurons treated with *Mecp2* KO ACM versus *Mecp2* WT ACM

| KO ACM vs. WT ACM | | | |
|---|---|---|---|
| Gene ID | Log2 Fold Change | P-value | FDR |
| Ostn | 0.6891 | 2.66E-04 | 0.0401 |
| Lamp5 | 0.5080 | 2.99E-04 | 0.0409 |
| Bdnf | 0.2198 | 1.64E-05 | 0.0092 |
| Adcyap1 | 0.2071 | 6.89E-06 | 0.0052 |
| Hapln1 | 0.2031 | 3.77E-04 | 0.0471 |
| Nr4a3 | 0.2021 | 1.93E-08 | 0.0001 |
| Vgf | 0.2002 | 2.47E-06 | 0.0031 |
| Lbh | 0.1878 | 1.68E-06 | 0.0031 |
| Zic1 | 0.1816 | 2.90E-04 | 0.0405 |
| Zfp804a | 0.1810 | 4.06E-05 | 0.0123 |
| Msi2 | 0.1792 | 1.91E-06 | 0.0031 |
| Nfat5 | 0.1754 | 1.08E-04 | 0.0219 |
| Olfm3 | 0.1734 | 2.29E-04 | 0.0371 |
| Dgkk | 0.1712 | 6.69E-05 | 0.0158 |
| Lgi2 | 0.1648 | 1.09E-07 | 0.0004 |
| Gad1 | 0.1613 | 1.74E-08 | 0.0001 |
| Fat3 | 0.1558 | 3.36E-06 | 0.0031 |
| Zdbf2 | 0.1546 | 1.79E-04 | 0.0314 |
| Ubash3b | 0.1536 | 4.14E-04 | 0.0495 |
| Slc32a1 | 0.1518 | 3.43E-04 | 0.0454 |
| Peg10 | 0.1503 | 3.25E-04 | 0.0435 |
| Klhl13 | 0.1486 | 1.39E-04 | 0.0256 |
| Rcan2 | 0.1467 | 2.83E-04 | 0.0405 |
| Cacna2d2 | 0.1456 | 2.63E-04 | 0.0401 |
| Kit | 0.1444 | 9.77E-05 | 0.0203 |
| Sipa1l2 | 0.1402 | 3.49E-04 | 0.0456 |
| Mindy2 | 0.1383 | 2.86E-04 | 0.0405 |
| Ccdc50 | 0.1377 | 2.81E-05 | 0.0107 |
| Hsd17b12 | 0.1353 | 3.36E-05 | 0.0117 |
| Dmxl1 | 0.1345 | 4.01E-05 | 0.0123 |
| Rab3c | 0.1331 | 3.02E-06 | 0.0031 |
| Slc6a1 | 0.1280 | 1.07E-05 | 0.0071 |
| Enah | 0.1275 | 2.90E-05 | 0.0107 |

| | | | |
|---|---|---|---|
| Fat1 | 0.1251 | 6.52E-05 | 0.0158 |
| Qk | 0.1243 | 2.82E-04 | 0.0405 |
| Gabra1 | 0.1243 | 3.95E-05 | 0.0123 |
| Fam126b | 0.1205 | 1.77E-04 | 0.0314 |
| Hdac9 | 0.1203 | 2.91E-05 | 0.0107 |
| Abat | 0.1183 | 2.76E-05 | 0.0107 |
| Rims1 | 0.1163 | 1.90E-05 | 0.0097 |
| Ttbk2 | 0.1140 | 3.97E-04 | 0.0483 |
| Dennd5b | 0.1119 | 2.31E-04 | 0.0371 |
| Herc3 | 0.1104 | 2.68E-04 | 0.0401 |
| Bcat1 | 0.1088 | 2.25E-04 | 0.0371 |
| Kcnh7 | 0.1080 | 3.94E-04 | 0.0483 |
| Chgb | 0.1039 | 2.92E-04 | 0.0405 |
| Cadps | 0.0997 | 5.53E-05 | 0.0148 |
| Gabrb1 | 0.0969 | 2.56E-04 | 0.0400 |
| Snhg11 | 0.0880 | 3.49E-05 | 0.0117 |
| Tuba1a | -0.0721 | 9.67E-05 | 0.0203 |
| Tubb5 | -0.0760 | 2.51E-04 | 0.0397 |
| Kif21b | -0.0898 | 3.69E-04 | 0.0469 |
| Gpm6b | -0.0904 | 3.16E-04 | 0.0428 |
| Ptms | -0.0956 | 3.57E-04 | 0.0461 |
| Atp1a1 | -0.0958 | 3.99E-04 | 0.0483 |
| Nell2 | -0.0976 | 1.09E-04 | 0.0219 |
| Ppp3ca | -0.0979 | 1.16E-04 | 0.0228 |
| Snca | -0.1043 | 2.20E-04 | 0.0368 |
| Camk2b | -0.1045 | 1.33E-05 | 0.0080 |
| Sptbn2 | -0.1094 | 1.88E-05 | 0.0097 |
| Hivep2 | -0.1155 | 6.32E-05 | 0.0158 |
| B3gat1 | -0.1210 | 1.23E-04 | 0.0239 |
| Acly | -0.1239 | 4.38E-05 | 0.0123 |
| Zbtb18 | -0.1326 | 2.33E-05 | 0.0107 |
| Ube2e3 | -0.1353 | 1.39E-04 | 0.0256 |
| Fam81a | -0.1357 | 1.83E-04 | 0.0314 |
| Atp2b4 | -0.1372 | 4.79E-05 | 0.0131 |
| Cdh13 | -0.1442 | 1.37E-06 | 0.0031 |
| Ildr2 | -0.1471 | 4.34E-05 | 0.0123 |
| Calb1 | -0.1475 | 1.61E-04 | 0.0291 |
| Wasf1 | -0.1493 | 2.32E-05 | 0.0107 |
| Sparcl1 | -0.1543 | 7.73E-06 | 0.0054 |

| | | | |
|---|---|---|---|
| Camk2a | -0.1544 | 2.48E-06 | 0.0031 |
| Rbfox3 | -0.1571 | 1.85E-04 | 0.0314 |
| Enc1 | -0.1572 | 5.65E-06 | 0.0045 |
| Camkk2 | -0.1623 | 2.59E-05 | 0.0107 |
| Cd24a | -0.1638 | 1.35E-05 | 0.0080 |
| Fabp5 | -0.1663 | 9.13E-05 | 0.0202 |
| Dtx4 | -0.1726 | 3.53E-05 | 0.0117 |
| Rfx3 | -0.1738 | 3.15E-06 | 0.0031 |
| Cpne4 | -0.1744 | 3.71E-04 | 0.0469 |
| Epha4 | -0.2043 | 1.53E-07 | 0.0004 |
| Nrgn | -0.2064 | 6.27E-05 | 0.0158 |
| Scd1 | -0.2126 | 6.75E-05 | 0.0158 |
| Dbpht2 | -0.2268 | 4.18E-05 | 0.0123 |
| Rnd1 | -0.2327 | 1.31E-04 | 0.0250 |
| Kcnv1 | -0.2417 | 6.34E-05 | 0.0158 |
| Cpne8 | -0.2427 | 2.54E-05 | 0.0107 |
| Ldb2 | -0.2470 | 5.40E-06 | 0.0045 |
| Ttc9b | -0.2480 | 8.11E-05 | 0.0186 |
| Wnt7b | -0.2556 | 2.96E-05 | 0.0107 |
| Igsf21 | -0.2643 | 9.32E-05 | 0.0202 |
| Sema4a | -0.2979 | 9.30E-05 | 0.0202 |
| Ipcef1 | -0.3034 | 2.90E-04 | 0.0405 |

**Supplemental Table 2:** Biological Processes GO Terms for Upregulated DEGs in wildtype neurons treated with *Mecp2* KO ACM versus *Mecp2* WT ACM samples

| Gene Ontology | Pathway | Enrichment FDR | Fold Enrichment | Genes |
|---|---|---|---|---|
| Biological Process | Gamma-Aminobutyric Acid Biosynthetic Process (GO:0009449) | 4.06E-03 | 224.1837 | Gad1 Abat |
| | Positive Regulation of Mast Cell Cytokine Production (GO:0032765) | 5.06E-03 | 179.3469 | Kit Nr4a3 |
| | Gamma-Aminobutyric Acid Import (GO:0051939) | 5.06E-03 | 179.3469 | Slc32a1 Slc6a1 |
| | Gamma-Aminobutyric Acid Metabolic Process (GO:0009448) | 7.14E-03 | 149.4558 | Abat Gad1 |
| | Negative Regulation of Synaptic Transmission GABAergic (GO:0032229) | 9.82E-03 | 112.0918 | Bdnf Slc6a1 |
| | Mast Cell Cytokine Production (GO:0032762) | 9.82E-03 | 112.0918 | Kit Nr4a3 |
| | Gamma-Aminobutyric Acid Transport (GO:0015812) | 3.32E-03 | 58.4827 | Slc32a1 Slc6a1 Abat |
| | Gamma-Aminobutyric Acid Signaling Pathway (GO:0007214) | 4.70E-03 | 46.3828 | Gabra1 Bdnf Gabrb1 |
| | Acidic Amino Acid Transport (GO:0015800) | 3.36E-03 | 24.5681 | Slc32a1 Slc6a1 Abat Bdnf |
| | Regulation of Postsynaptic Membrane Potential (GO:0060078) | 3.36E-03 | 14.7489 | Gabra1 Adcyap1 Gabrb1 Abat Bdnf |
| | Sensory Organ Morphogenesis (GO:0090596) | 5.06E-03 | 9.2130 | Bdnf Olfm3 Fat3 Nr4a3 Fat1 Zic1 |
| | Chemical Synaptic Transmission (GO:0007268) | 1.50E-03 | 6.2447 | Gabra1 Gabrb1 Vgf Bdnf Cadps Kit Adcyap1 Slc6a1 Abat Cacna2d2 |
| | Anterograde Trans-Synaptic Signaling (GO:0098916) | 1.50E-03 | 6.2447 | Gabra1 Gabrb1 Vgf Bdnf Cadps Kit Adcyap1 Slc6a1 Abat Cacna2d2 |
| | Trans-Synaptic Signaling (GO:0099537) | 1.50E-03 | 6.1759 | Gabra1 Gabrb1 Vgf Bdnf Cadps Kit Adcyap1 Slc6a1 Abat Cacna2d2 |
| | Behavior (GO:0007610) | 1.50E-03 | 6.0508 | Bdnf Kit Adcyap1 Slc6a1 Vgf Abat Rcan2 Nr4a3 Zic1 Gad1 |

| | | | | |
|---|---|---|---|---|
| | Synaptic Signaling (GO:0099536) | 1.52E-03 | 5.9151 | Gabra1 Gabrb1 Vgf Bdnf Cadps Kit Adcyap1 Slc6a1 Abat Cacna2d2 |
| | Secretion (GO:0046903) | 9.24E-03 | 4.2140 | Rab3c Cadps Adcyap1 Kit Slc6a1 Rims1 Abat Vgf Nr4a3 Bdnf |
| | Nervous System Development (GO:0007399) | 1.60E-04 | 3.6542 | Hapln1 Enah Adcyap1 Zic1 Lgi2 Bdnf Gabra1 Hdac9 Kit Olfm3 Gabrb1 Slc32a1 Rims1 Ostn Abat Zfp804a Nr4a3 Qk Fat3 Ttbk2 |
| | System Development (GO:0048731) | 1.66E-03 | 2.4090 | Kit Hapln1 Enah Adcyap1 Ubash3b Zic1 Lgi2 Bdnf Gabra1 Ostn Hdac9 Olfm3 Gabrb1 Slc32a1 Rims1 Abat Zfp804a Fat3 Nr4a3 Vgf Qk Fat1 Ttbk2 Peg10 |
| | Multicellular Organism Development (GO:0007275) | 2.46E-03 | 2.2672 | Kit Hapln1 Enah Adcyap1 Zdbf2 Ubash3b Zic1 Lgi2 Bdnf Ostn Gabra1 Hdac9 Olfm3 Gabrb1 Slc32a1 Rims1 Abat Zfp804a Fat3 Nr4a3 Vgf Qk Fat1 Ttbk2 Peg10 |

**Supplemental Table 3:** Biological Processes GO Terms for Downregulated DEGs in wildtype neurons treated with *Mecp2* KO ACM versus *Mecp2* WT ACM samples

| Gene Ontology | Pathway | Enrichment FDR | Fold Enrichment | Genes |
|---|---|---|---|---|
| Biological Process | Negative Regulation of Neurotransmitter Uptake (GO:0051581) | 2.52E-03 | 249.6591 | Gpm6b Snca |
| | Negative Regulation of Serotonin Uptake (GO:0051612) | 2.52E-03 | 249.6591 | Gpm6b Snca |
| | Long-Term Synaptic Potentiation (GO:0060291) | 4.25E-04 | 24.0057 | Calb1 Nrgn Epha4 Snca Camk2b |
| | Regulation of Calcium-Mediated Signaling (GO:0050848) | 2.52E-03 | 23.7771 | Ppp3ca Cd24a Atp2b4 Cdh13 |
| | Regulation of Synapse Organization (GO:0050807) | 2.77E-04 | 13.8151 | Tubb5 Tuba1a Snca Epha4 Sema4a Sparcl1 Camk2b |
| | Regulation of Synaptic Plasticity (GO:0048167) | 8.92E-04 | 13.4951 | Calb1 Nrgn Epha4 Camk2a Snca Camk2b |
| | Regulation of Synapse Structure or Activity (GO:0050803) | 2.77E-04 | 13.4432 | Tubb5 Tuba1a Snca Epha4 Sema4a Sparcl1 Camk2b |
| | Synapse Organization (GO:0050808) | 4.43E-06 | 11.3481 | Epha4 Snca Tubb5 Wasf1 Sparcl1 Sptbn2 Tuba1a Camk2b Wnt7b Sema4a Igsf21 |
| | Cell Junction Organization (GO:0034330) | 1.30E-05 | 7.9573 | Epha4 Snca Tubb5 Wasf1 Sparcl1 Sptbn2 Tuba1a Gpm6b Camk2b Wnt7b Sema4a Igsf21 |
| | Regulation of Trans-Synaptic Signaling (GO:0099177) | 1.08E-03 | 7.8478 | Calb1 Snca Fabp5 Nrgn Epha4 Camk2a Ppp3ca Camk2b |
| | Trans-Synaptic Signaling (GO:0099537) | 3.44E-04 | 6.8777 | Snca Calb1 Camk2a Fabp5 Nrgn Epha4 Ppp3ca Cd24a Camk2b Sptbn2 |
| | Synaptic Signaling (GO:0099536) | 4.25E-04 | 6.5873 | Snca Calb1 Camk2a Fabp5 Nrgn Epha4 Ppp3ca Cd24a Camk2b Sptbn2 |
| | Chemical Synaptic Transmission (GO:0007268) | 1.35E-03 | 6.2589 | Snca Calb1 Camk2a Nrgn Epha4 Ppp3ca Cd24a Camk2b Sptbn2 |

| | | | | |
|---|---|---|---|---|
| | Anterograde Trans-Synaptic Signaling (GO:0098916) | 1.35E-03 | 6.2589 | Snca Calb1 Camk2a Nrgn Epha4 Ppp3ca Cd24a Camk2b Sptbn2 |
| | Neuron Development (GO:0048666) | 6.41E-04 | 4.8993 | Epha4 Sema4a Gpm6b Wnt7b Cd24a Camk2a Enc1 Rnd1 Wasf1 Ppp3ca Zbtb18 Camk2b |
| | Cell-Cell Signaling (GO:0007267) | 2.90E-04 | 4.5570 | Wnt7b Snca Calb1 Camk2a Fabp5 Ppp3ca Nrgn Epha4 Cd24a Ildr2 Rfx3 Camk2b Nell2 Sptbn2 |
| | Nervous System Development (GO:0007399) | 3.55E-05 | 3.8660 | Wnt7b Rbfox3 Epha4 Sema4a Gpm6b Ldb2 Cd24a Camk2a Snca Atp2b4 Ppp3ca Enc1 Nrgn Rnd1 Camk2b Wasf1 Igsf21 Zbtb18 Sptbn2 |
| | Movement of Cell or Subcellular Component (GO:0006928) | 1.35E-03 | 3.4917 | Epha4 Sema4a Rnd1 Wnt7b Wasf1 Camk2a Atp2b4 Ppp3ca Cdh13 Kif21b Camk2b Ldb2 Rfx3 Cd24a Atp1a1 |
| | Regulation of Biological Quality (GO:0065008) | 7.03E-06 | 3.1991 | Atp2b4 Sema4a Calb1 Gpm6b Atp1a1 Rnd1 Ppp3ca Tubb5 Camk2a Snca Tuba1a Epha4 Nrgn Sptbn2 Fabp5 Ldb2 Wnt7b Scd1 Ildr2 Rfx3 Cd24a Camk2b Zbtb18 Nell2 Sparcl1 |
| | System Development (GO:0048731) | 1.35E-03 | 2.4591 | Wnt7b Rbfox3 Epha4 Sema4a Gpm6b Cdh13 Ldb2 Cd24a Camk2a Snca Atp2b4 Ppp3ca Enc1 Nrgn Rnd1 Camk2b Calb1 Wasf1 Rfx3 Igsf21 Zbtb18 Sptbn2 |
